# Correlating Protein Aggregate Structure with Cellular Function in Differentiated Muscle Cells: Discriminating Pathogenic from Non-Pathogenic Forms

**DOI:** 10.1101/2024.04.25.591067

**Authors:** Sander D. Mallon, Erik Bos, Vahid Sheikhhassani, Milad Shademan, Dino Rocca, Lenard M. Voortman, Alireza Mashaghi, Thomas H. Sharp, Vered Raz

## Abstract

Ageing has a major adverse impact on maintaining cellular proteostasis and age-related dysregulation leads to an increase in protein aggregation. Equivalently, the accumulation of aggregated proteins accelerates proteostasis impairment. Accumulation of protein aggregates and impaired proteostasis are hallmarks of ageing-associated neuromuscular disorders and tissue degeneration is predominantly in post-mitotic muscle and neuronal cells. A short alanine expansion mutation in the Poly(A) binding protein nuclear 1 (PABPN1) causes Oculopharyngeal muscular dystrophy (OPMD), a rare age-associated protein aggregation myopathy. PABPN1 is a vital RNA-binding protein but OPMD pathology is limited to skeletal muscles connected to nuclear aggregates. In contrast to the mutant PABPN1, the wild-type PABPN1 forms age-associated non-pathogenic aggregates.

We generated an inducible muscle cell models for mutant and wild-type PABPN1 protein aggregation. By combining four different, but complementary, imaging modalities, covering micro- to nanoscale resolutions, we were able to characterise differences in structure and dynamics between pathogenic and non-pathogenic PABPN1 aggregates in differentiated muscle cells. These data allowed us to correlate the structure of aggregates to cellular function, providing important insights into how aggregates lead to cell dysfunction in post-mitotic cells.

**Graphical summary:** 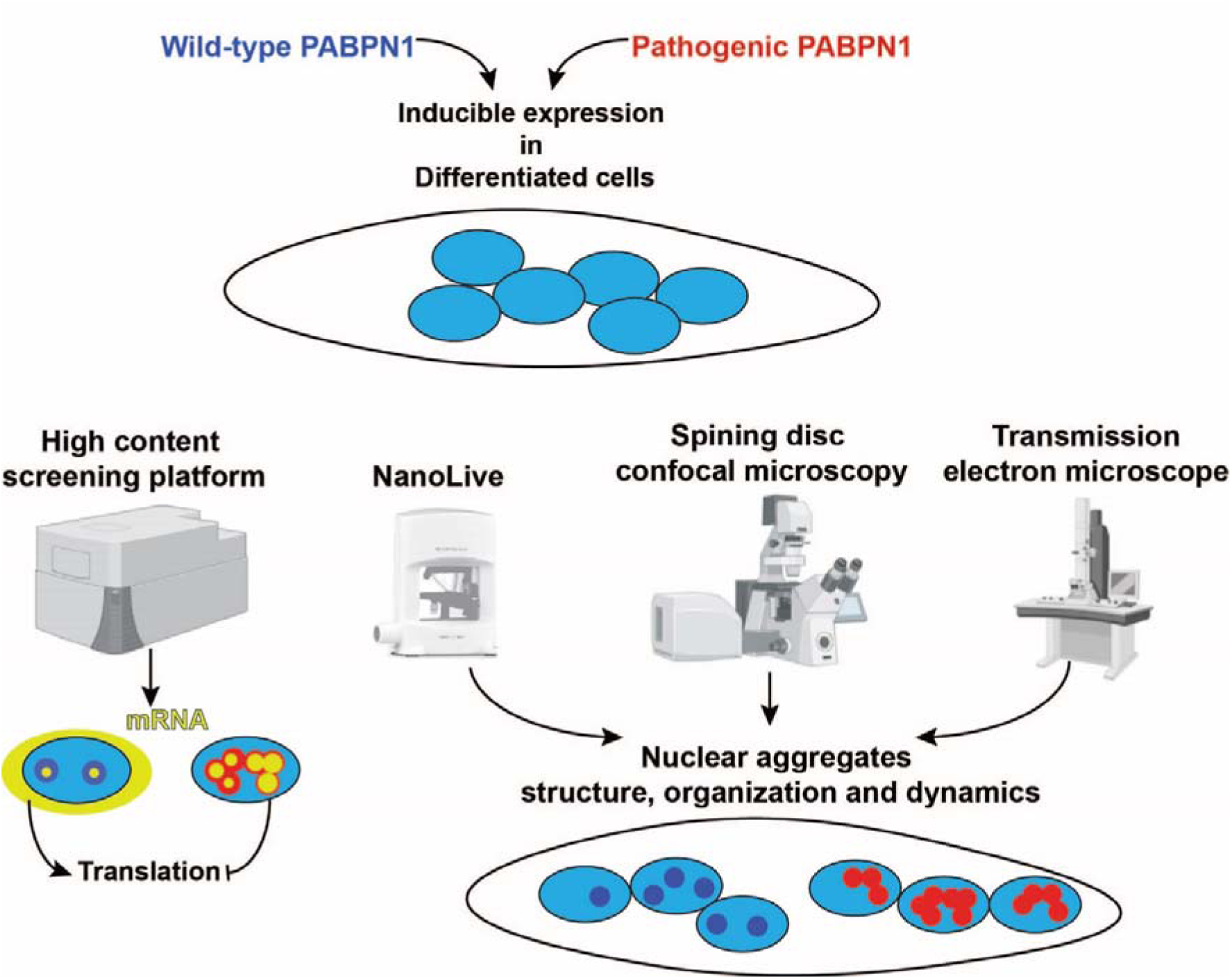

## Introduction

Maintaining protein homeostasis, or proteostasis, is a fundamental aspect of cellular function, particularly crucial for neurons and muscle cells (Hipp, Kasturi and Hartl 2019). Proteostasis encompasses a complex network of pathways that govern the synthesis, folding, trafficking, and degradation of proteins within the cell. This network ensures that proteins maintain their natively-folded functional conformations, preventing protein misfolding and aggregation that can lead to cellular dysfunction and disease (Scalvini, Heling et al. 2023, Wruck, Avellaneda et al. 2023). A balance in proteostasis is particularly crucial in post-mitotic cells, such as differentiated neurons and skeletal muscle fibers. In neuronal and skeletal muscle tissues, proteostasis imbalance leads to protein aggregates that adopt a similar and highly ordered insoluble amyloid structure (Aguzzi and O’Connor 2010, McCormick and Vasilaki 2018). The maintenance of proteostasis is a delicate balance that is prone to disruption by ageing-associated stresses leading to cell dysfunction and tissue degeneration (Taylor and Dillin 2011).

Ageing is a primary factor that challenges the integrity of the proteostasis machinery (Cuanalo-Contreras, Schulz et al. 2022, Frankowska, Lisowska and Witkowski 2022). Over time, the efficiency of proteostatic mechanisms declines, leading to an increase in protein misfolding and aggregation. The relationship between ageing and proteostasis is bidirectional: not only does ageing contribute to proteostasis decline, but impairment in proteostasis can also accelerate the ageing-associated tissue degeneration and the onset of associated diseases (Morimoto and Cuervo 2014). This complex interplay underscores the importance of understanding proteostasis in the context of ageing and disease, and to develop interventions that could promote healthy ageing and mitigate disease progression. Although age-associated decline in cell function affects most cell types, many diseases of the neuromuscular system are associated with the accumulation of insoluble protein aggregates. The pathogenic protein aggregates can be cytoplasmic, such as in Parkinson’s disease and Alzheimer’s disease, or nuclear as found in polyglutamine repeat disorders, including Huntington’s disease (HD) and Spinocerebellar ataxia diseases (SCAs), or polyalanine repeat disorders such as Oculopharyngeal muscular dystrophy (OPMD) (Abu-Baker and Rouleau 2006).

OPMD is caused by a short expansion mutation in the gene that encodes poly(A)-binding protein nuclear 1 (PABPN1) (Brais, Bouchard et al. 1998). The wild type PABPN1 has a repeat of 10 alanine residues at the N-terminus, but the expansion results in the mutant protein containing 11-18 alanine expansion. PABPN1 is ubiquitously expressed and essential in all eukaryotic cells, but symptoms are limited to skeletal muscles. Both wild type and the expanded PABPN1 are prone to aggregation, but only aggregates of the mutant protein are pathogenic (Yamashita 2021). Discriminating between structural features of the pathogenic PABPN1 aggregates from the non-pathogenic form has been addressed in previous studies utilizing mitotic, non-muscle cells (Raz, Abraham et al. 2011, Guan, Jiang et al. 2023). However, these cell models do not fully capture the nuances of proteostasis in post-mitotic cells or muscle-specific proteostasis (Bar-Lavan, Shemesh et al. 2016). Here, we aim to provide this information by examining the structural differences between pathogenic and non-pathogenic PABPN1 alleles within a human muscle cell model. We generated stable muscle cells expressing the wild-type PABPN1 (Ala10) or the pathogenic allele (Ala16) under the tetracycline-inducible promoter. The inducible promoter allows investigation of the aggregated PABPN1 in differentiating muscle cells and bypasses PABPN1 effect in proliferating cells. We investigated the structure of aggregated PABPN1 with five imaging modalities, ranging from micrometer to nanometer resolution in living and fixed differentiated muscle cells. Detailed image analysis enabled quantification of structural differences between Ala10 and Ala16 proteins that are associated with cellular processes of muscle cells and PABPN1 function.

## Methods

### PABPN1 constructs and lentivirus production

The full length PABPN1 wild type (Ala10) or the expanded PABPN1 (Ala16) fused to YFP, previously described in (Raz, Abraham et al. 2011) were cloned into the pCW57-MCS1-2A-MCS2 doxycycline (Dox) inducible lentiviral vector (Addgene plasmid #71782). Cloning was confirmed by sanger sequencing. Lentivirus production was performed as detailed in (Carlotti, Bazuine et al. 2004). Lentivirus particle titers were determined in HeLa cells.

### Cell culture

Cells were cultured in growth medium (F10 (Gibco) medium supplemented with 15% FCS, 1Lng/ml bFGF, 10Lng/ml EGF and 0.4Lµg/ml Dexamethasone). Cells were propagated in confluence 50-80%. Cell cultures did not reach 100% confluence to avoid spontaneous differentiation. Cell differentiation was induced at high confluency (85-95%) in DMEM+2% horse serum for 3-5 days. The 2417 immortal human muscle cells were transduced with lentiviruses encoding Ala10-YFP or Ala16-YFP, and stable cell cultures were created using puromycin selection. Induction of the PABPN1-YFP transgene was carried out with 4 μg/ml doxycycline hyclate (D5207, Sigma Aldrich). For high content screening (HCS), cells were seeded in a Nunc 96 well plate; for live cell confocal microscopy, cells were seeded in either a µ-Slide 15 Well 3D ibiTreat (81506, IBIDI) or a µ-Slide 8 Well high ibiTreat (80806, IBIDI) slide. For electron microscopy, cells were seeded in µ-Dish 35 mm, high Grid-500 ibiTreat (81166, IBIDI) dishes. For refractive index imaging, cells were cultured on a µ-Dish 35 mm, low ibiTreat (80136, IBIDI) dish. Cell cultures were treated with 0.5 µM epoxomicin (Sigma-Aldrich #134381-21-8) for 6 hours; Leptomycin B (LMB) 20nM for 4 hours; or 20 µg/ml cycloheximide (Sigma-Aldrich # 01810) for one hour. Cells were incubated in the growth medium.

### Protein extraction and Western blotting

Proteins were lysed from cells using RIPA buffer (20 mM Tris, pH 7.4, 150 mM NaCl, 5 mM EDTA, 1% NP40, 5% glycerol and 1 mM DTT and protease inhibitor cocktail). After sonication and centrifugation (1 min, 13000g, at 4°C), the supernatant containing the soluble proteins was transferred to a new tube and the pellet, containing the insoluble proteins, was washed once in PBS, dissolved in loading buffer, sonicated and spin down prior to heat inactivation. Protein aliquots were separated on 10% SDS-PAGE. Western blotting was carried out with a PVDF membrane. Bulk proteins were visualized with the No-Stain Protein Labeling Reagent (#A44717, ThermoFisher) and imaged using the iBright Imaging System (ThermoFisher). The membrane was blocked with 5% dried milk powder (T145.2, Carl Roth), first antibody incubation was carried out at 4 degrees overnight, and secondary antibody incubation at room temperature for one hour. Antibodies are listed in Table S1. An Odyssey CLx Infrared imaging system (LiCOR, NE. USA) was used to detect the fluorescent signal. Quantification of protein abundance was done using ImageJ. Values were corrected for background and normalized to loading controls. Western blot quantification was carried out with ImageJ. Normalization was made for both the No-Stain and house-keeping signal.

### Immunohistochemistry

Insoluble PABPN1 was detected in 1M KCl pre-treatment for 15 minutes. Immunohistochemistry with or without KCl pre-treatment was performed using standard procedures: fixation (2% formaldehyde in PBS) for 5 minutes, permeabilization (1% triton in PBS) for 10 minutes, PBS washing, incubation with primary antibodies for one hour at room temperature, PBS washing, incubation with secondary antibodies +DAPI (4L,6-Diamidino-2-phenylindole dihydrochloride) for 30 minutes, and PBS washing. Cells were kept in PBS during imaging. Antibodies are listed in Table S1.

### Cellular Assays

The Mitochondrial activity assays were performed in differentiated or proliferating cell cultures grown in 96 well plate and treated with Tetramethylrhodamine methyl ester perchlorate (TMRM). Cell cultures grown in 96 well plates were washed with PBS and incubated with a staining solution (5nM TMRM (Sigma-Aldrich #115532-50-8) and Hoechst were diluted in growth media and incubated for 45 minutes. After twice PBS washing, cells were kept in differentiation media during imaging.

The Protein Synthesis Assays were performed in differentiated or proliferating cell cultures grown in 96 well plate. The protein synthesis assay kit (Cayman Chemicals #601100) was employed according to the manufacturer protocol, with the following modifications: azido-*O*-propargyl-puromycin (OPP)- 488 was replaced with OPP-555 (Vector laboratories, #CCT-1494). For the negative control, 30 minutes pre incubation with 20µM cycloheximide was used. Hoechst was added after fixation.

For RNA hybridization with Oligo-dT, differentiated cell cultures were fixed using 3.7% FA for 15 minutes at RT. After two PBS washes the cells were incubated in Protease III diluted 1:30 in PBS (#322337 Advanced Cell Diagnostics) for 15 minutes at RT. After twice PBS washes cells were incubated in hybridization buffer (#10369 Cepham Life Sciences) for 15 minutes at RT. Incubation with 5’-Cy5-Oligo d(T)12-18 probe (#26-4400-02 Gene Link), diluted 1:1000 in hybridization buffer, was carried out overnight at 40 degrees in a humidified chamber. The following day, washes were carried out at 40 degrees for 5 minutes with 4x, 2x, 1x SSC buffer, and with PBS. Finally, the cells were incubated with Hoechst and kept in PBS during imaging.

For differentiation index calculation, cells in 96 well plate were treated with Dox for 24 hours, and then were incubated in differentiation medium for 72 hours. Differentiated cells were marked for MyHC expression using Immunohistochemistry procedure and the MF20 antibody.

### Imaging and Image quantification

The CellInsight CX7 LZR high content screening (HCS) platform was used for high-content imaging. A cell-based analysis was performed with the accompanied HCS Platform spot detector and co-localization toolbox (ThermoFisher Scientific). Cells were imaged with 405 nm (DAPI), 488 nm (YFP) filters, and per cellular assay with the following filters: imaging: TMRM with 560 nm, OPP-555 with 560 nm, oligo-dT-Cy5 with 647 nm, and MF20 antibody with 647 nm.

Imaging for calculation of differentiation index was made with a 10x objective covering over 12,000 nuclei per well, and imaging for nuclear YFP quantification with a 20x objective, covering at least 5000 nuclei per well. The co-localization toolbox was used for the quantification of differentiation index by the percentage of myonuclei without MyHC objects, and the spot detection toolbox for the YFP puncta, TMRM, OPP-555 and oligo-dT-Cy5. Bulk YFP intensity was considered with a low threshold (50-150) and the high threshold with 650-1200. The exact threshold was adjusted per experiment, per experiment both low and high thresholds were quantified. Analysis of both TMRM and 555-OPP MFI was made from the perinuclear region. Oligo-dT signal and overlap with YFP was measured from both nuclear and perinuclear regions in YFP positive myonuclei. The calculation of the average MFI, spot, nuclear/perinuclear MFI ratio or colocalization was per nucleus over the entire well.

Confocal microscopy imaging: imaging of fixed single nuclei (in Fig. 2) was made with Leica SP8 confocal microscope using a 63 x/1.3 oil objective and HyD detectors. DAPI nuclear counterstain was imaged with 405 nm and YFP with white light laser at 488 nm. All other experiments, using confocal microscopy, were images with a Leica DMi8 with the Andor Dragonfly spinning disc module using a 40x/1.3 or 63x/1.3 oil immersion objective. Nuclear counterstain DAPI or SiR-700 DNA (Cy-SC015, Cytoskeleton, Inc.) was imaged with a laser at 405 nm or 647 nm, respectively, the YFP signal at 488 nm and oligo-dT at 647 nm. Identical imaging settings including exposure time, laser power, the excitation-emission range, and Z-stacks step size were employed within an experiment. Time-lapse imaging was made with Z-stack acquisition taking 4.20 minutes (1 frame per 2 seconds). Maximum-projection of Z-stacks was made with Imaris.

Quantifications of confocal images were carried out in ImageJ.

- Overlay of time-lapse images was made from frames at 0, 130, and 260 seconds. Each frame was assigned a distinct RGB color, and an overlap was made with imageJ.

*-* YFP puncta quantification was carried out with a macro in ImageJ. The YFP channel was processed with a gaussian blur (1.5 sigma (radius) ‘Blur’ and ‘Despeckle’ functions. Masking of the YFP puncta was applied with a constant threshold across all images within an experiment. The threshold was manually determined to match the YFP puncta (examples are in Fig. S3). Particles >0.01 μm^2^ in area were considered for analysis. From each punctum, the mean fluorescence intensity (MFI), the area and the circularity were recorded. Analysis was carried out on gated myonuclei from fused cells or unfused cells (single nucleus).

-Oligo-dT signal analysis was carried out on gated myonuclei in fused cells in ImageJ. YFP puncta and oligo-dT analysis was carried out with a Gaussian blur of 1.0, per fluorophore, a constant threshold was used for all images. The MFI and area were recorded. The overlap and correlation between YFP and oligo-dT were assessed with the JACoP plugin (Bolte and Cordelieres 2006) in ImageJ, using the M1 & M2 coefficients and the Pearson correlation.

### Electron microscopy

Differentiated cell cultures were fixed in 1.5% glutaraldehyde in 0.1 M Sodium Cacodylate buffer for 2 hours and were successively incubated in 1% Osmium Tetroxide in 0.1 M cacodylate buffer for 1 hour and in 1% Uranyl Acetate in water for 1 hour. The cells were then dehydrated through a series of incubations in Ethanol (70-100%) for 90 minutes and embedded in Epon. The flat embedded cells were sectioned with an ultramicrotome (UC6, Leica, Vienna) using a 35 degrees diamond knife (Diatome, Biel, Switzerland) at a nominal section thickness of 90 nm. The sections were transferred to a formvar and carbon coated 1×2 mm copper slot grid and stained for 20 minutes with 7% uranyl acetate in water and for 10 minutes with lead citrate. EM images were recorded using a Tecnai 12 electron microscope (Thermo Fisher Scientific) equipped with an EAGLE 4k×4k digital camera. For navigation on EM images, montages of images at 11,000× were generated using stitching software (Faas, Avramut et al. 2012). Morphology was assessed by sampling 100 nuclei on stitched EM images. For 3D reconstruction, 8 consecutive serial sections with a nominal thickness of 200 nm were manually aligned, segmented using the software program, Ais (Last, Abendstein et al. 2024) and rendered as 3D isosurfaces in ChimeraX (Pettersen, Goddard et al. 2021). For correlative light and electron microscopy, cells were grown on a gridded µ-Dish 35 mm plate and stained with DAPI. The living cells were imaged in an EVOS FL Digital Inverted Fluorescence Microscope (Invitrogen) equipped with a 20× objective. YFP was visualised with a GFP filter and DAPI with a UV filter. After imaging in the light microscope, the cells were processed for electron microscopy as described above. Superimposition and correlation of light and electron microscopy images was performed using Photoshop. In Photoshop the LM image was copied as a layer into the EM image and made 50% transparent. The LM image required transformation to align with the broader scale of the EM image. This involved isotropic scaling and rotation. Alignment was facilitated by utilizing the nuclear DAPI staining alongside cell morphology.

### Holo-tomographic microscopy (HTM)

HTM, in combination with epifluorescence, was performed on the 3D Cell-Explorer Fluo (Nanolive, Ecublens, Switzerland) using a 60× air objective (NA = 0.8) at a wavelength of λ = 520 nm (Class 1 low power laser, sample exposure 0.2 mW/mm2) and CMOS Sony sensor, with quantum efficiency (typical) 70% (at 545 nm), dark noise (typical) 6.6 e-, dynamic range (typical) 73.7 dB, field of view 90 × 90 × 30 μm, axial resolution 400 nm, and maximum temporal resolution 0.5 3D RI volume per second. Acquired RI images were processed with built-in software (Nanolive). ImageJ/Fiji (https://imagej.nih.gov/) was used for the final processing and quantifications.

### Nuclei texture analysis

RI images of each cell type were initially converted to the 8-bit format using ImageJ. Subsequently, the areas, including the cell nuclei, were outlined, and extracted using a free-hand selection tool. Following this, the texture of these selected areas was analyzed utilizing the GLCM (Gray Level Co-occurrence Matrix) texture analysis plug-in (version 0.4) developed by Julio E. Cabrera. This plug-in facilitated the computation of various statistical parameters associated with the GLCM of the image, including the Inverse Difference Moment (IDM) and Entropy.

### Statistics

Statistical tests were performed in GraphPad Prism.

## Results

### Ala16-YFP aggregation is higher than Ala10-YFP in an inducible muscle cell model

We generated human muscle cells that stably express the Ala10 wildtype PABPN1 (named here Ala10) or the Ala16 expanded PABPN1 (named here Ala16); both were fused to YFP and under the tetracycline-inducible promoter. Stable cells were made by lentivirus transduction that ensure low copy number of the transgene. We first assessed PABPN1 aggregation using Western blot analysis. The PABPN1-YFP protein was not detected in uninduced (vehicle) cells, but after Doxycycline (Dox) treatment both Ala10 and Ala16 proteins were detected (Fig. 1A). A prominent accumulation of both Ala10 and Ala16 proteins was found in the insoluble fraction, but the Ala16 accumulation in the insoluble fraction was higher compared with Ala10 (Fig. 1B). This observation contrasts with studies in non-muscle proliferating cells showing limited or no difference between insoluble levels of the wild type and the pathogenic PABPN1 (Guan, Jiang et al. 2023). Next, we assessed whether the transgene overexpression is associated with the endogenous PABPN1 levels and found a higher ratio in Ala16 compared with Ala10 or in vehicle cultures (Fig. 1C). This suggests that an increase in the insoluble PABPN1 depletes levels of the soluble form.

**Figure 1.**
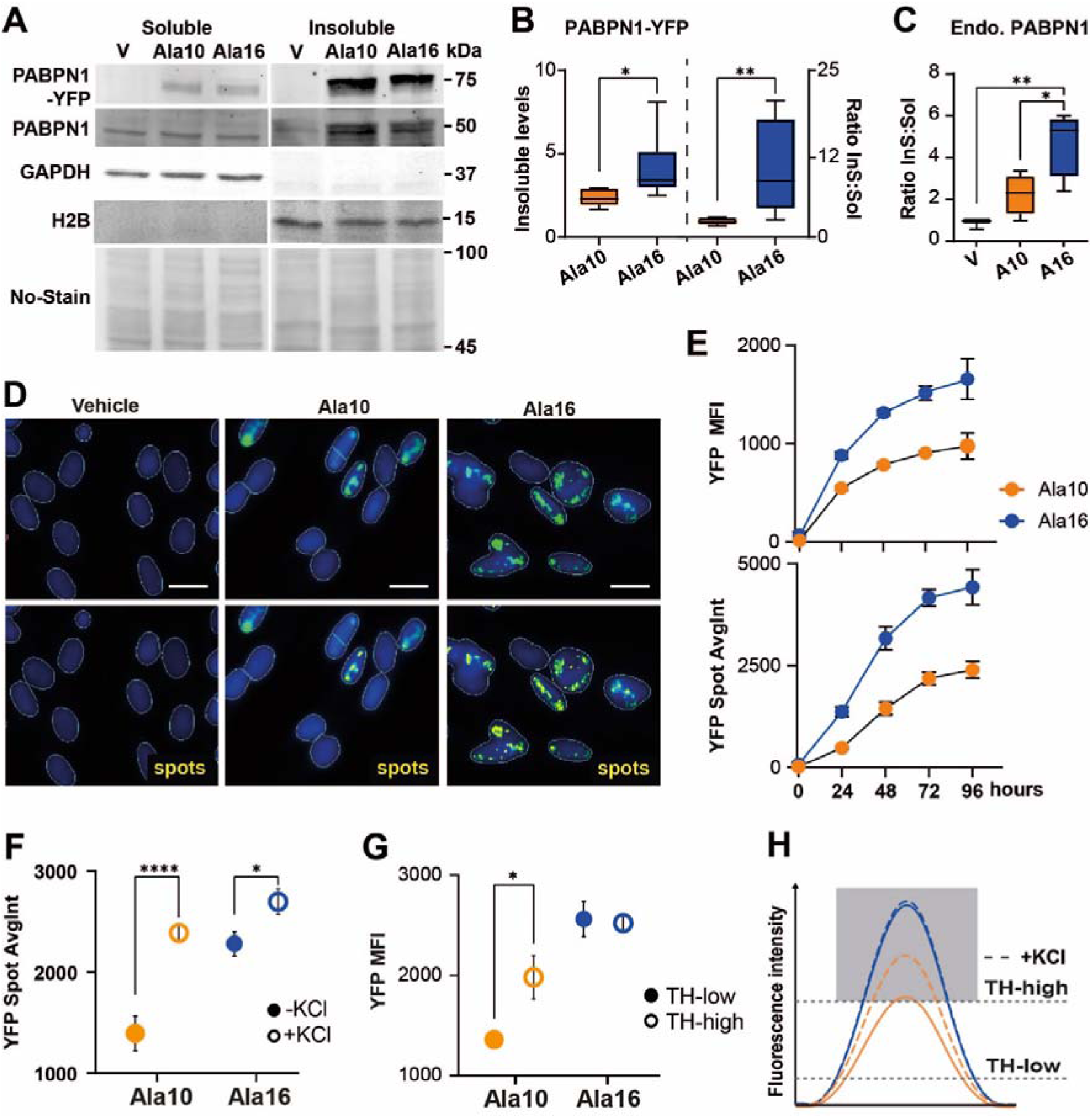
Aggregation characterization of Ala10 and Ala16 in muscle cells. Panels (A-G) were carried out in proliferating cell cultures incubated with Dox for 72 hours **A.** A Western blot of the soluble (sol) and insoluble (InS) fractions from vehicle (V) Ala10-YFP and Ala16-YFP cell cultures. The PABPN1-YFP is 75 kDa, the endogenous PABPN1 is 50 kDa, GAPDH or H2B is control for soluble or insoluble fractionations, respectively. The No-Stain is a loading control. **B.** Boxplot of the insoluble PABPN1-YFP levels in Ala10 and Ala16 (left) or the insoluble/soluble ratio (right). Expression levels were normalized to the No-stain and the Ala10 soluble fraction. The mean and standard deviation are from N=6. or **C.** Boxplot of the endogenous (Endo.) The PABPN1 insoluble/soluble ratio in vehicle (V), Ala10, and Ala16. Expression levels were normalized to the No-stain. The mean and standard deviation are from N=4. **D.** Representative images of YFP (green) and the segmented puncta in yellow (spots) in vehicle, Ala10 and Ala16 myonuclei. Scale bar is 10μm. **E.** Dot plot shows bulk YFP average intensity in Ala10 and Ala16 myonuclei with and without KCl treatment. **G.** Dot plot shows YFP intensity at low or high threshold. **H.** A schematic presentation of intensity of Ala10-YFP (orange) or Ala16-YFP (blue) after KCl treatment (dashed curved line), or at low or high YFP threshold. The low threshold measures bulk YFP intensity and the high threshold the aggregated signal. The grey area presents the aggregated puncta. **H.** Accumulation of bulk YFP intensity (lower panel) and high threshold of YFP intensity (right panel) over time in Ala10 or Ala16 cells. Average and standard deviation are from N=3 biological replicates. Statistical significance was assessed with a t-test parametric test. P<0.05 or <0.0001 is denoted with *, ****, respectively.

**Figure 2.**
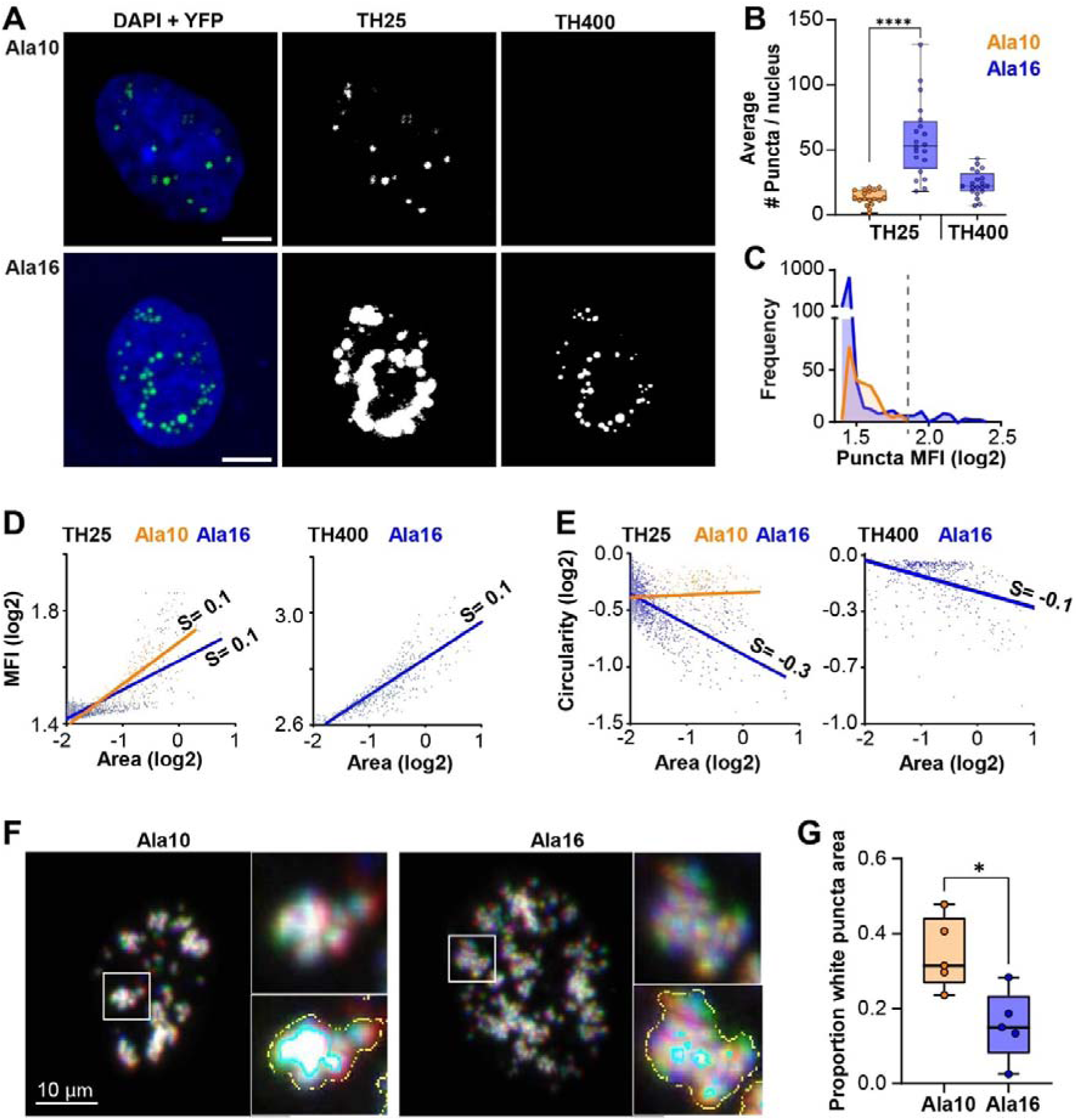
Structural features of Ala10 and Ala16 aggregates. Experiments in panels A-E were carried out in differentiated cell cultures with 72 hours Dox treatment. 20 nuclei per genotype were used for analysis (B-E). **A.** A representative confocal image of Ala10-YFP and Ala16-YFP (in green) in a single myonuclei (DAPI counter stain is in blue), and masks of the YFP puncta at low and high threshold (TH25, and TH400, respectively). The YFP in the fluorescence image was differentially stretched for visualization purpose. Scale bar is 10μm. **B.** A bar chart shows the average puncta number per nucleus in Ala10 and Ala16 myonuclei. Average and standard deviation are from Statistical significance was determined by an ordinary one-way ANOVA test. P<0.0001 is denoted with ****. C. Frequency distribution plot of Ala10 (orange) and Ala16 (blue) puncta mean fluorescence intensity (MFI) (log2). The dashed line depicts the cutoff (1.3-1.85 log2 MFI) that was used for Ala10 and Ala16 puncta comparison (D-E). D-E. Scatter plots of puncta area (log2) vs. MFI (log2) (D), or puncta area (log2) vs. Circularity (log 2) (E) in TH25 and TH400. The linear regression line is depicted per genotype, and the significant slope (S) is denoted. F. Proliferating cell cultures were imaged 16 hours after Dox treatment. Puncta dynamics is shown by the overlay of three-time frames. Each time is depicted in red, green or blue, a puncta overlay is depicted in white. A magnification of the boxed area is and the segmented area of white color (threshold 225 in cyan line) and all colors (threshol 90 in yellow line). G. A bar chart shows the proportion of white area to colour area in Ala10 and Ala16 nucleus. Average and standard deviation are from Statistical significance was determined by the Student’s test. P<0.05 is denoted with *.

Fluorescence imaging confirmed nuclear accumulation on both Ala10 and Ala16 (Fig. 1D). Consistent with the Western blot, the YFP fluorescence intensity in Ala16 was higher than in Ala10 (Fig. 1E). The aggregated YFP was monitored after KCl treatment (representative images are in Fig. S1), which clears the insoluble proteins and intensifies the insoluble puncta, resulting in higher YFP intensity after KCl treatment (Fig. 1E). A larger difference was found in Ala10 compared to Ala16 (Fig. 1E), indicating that the YFP signal in the Ala10 cells is partly of the soluble protein and partly of the aggregated protein, whereas in Ala16 the signal is predominantly of the aggregated protein. To identify PABPN1 aggregates without KCl treatment, we applied different thresholds that differentiate between the signal of the aggregated protein from the bulk nuclear YFP signal, and the results were consistent with the KCl treatment (Fig. 1F). A high threshold gave similar results as KCl treatment (Fig. 1F), as schematically summarized in Fig. 1G.

We then applied two thresholds to discriminate between bulk YFP accumulation over time and aggregation. Bulk YFP accumulation showed a similar curve in Ala10 and Ala16 and a plateau was reached after 48 hours (Fig. 1H). The puncta fluorescence intensity reached a plateau after 72 (Fig. 1H), confirming that aggregation is delayed compared to expression.

### Structural features of PABPN1 aggregates

To study aggregation in differentiated muscle cells, we focused on multinucleated cells which can be recognized by the expression of the myosin heavy chain (Fig. S2A). Overexpression of Ala10 or Ala16 prior to differentiation resulted in reduced differentiation index regardless of the genotype (Fig. S2B). Therefore, in all experiments, cell cultures were treated with Dox during differentiation and transgene expression was initiated after induction of differentiation. The fully differentiated cells can also be recognized by myonuclei clustering, indicating cell fusion (Fig. S2C). We used myonuclei clusters to identify fused cells.

To identify structural features, single nuclei were imaged with a confocal microscope (Fig. 2A), and YFP puncta were segmented with a constant, manually selected, threshold (Fig. S3). Unlike the HCS image quantification, with confocal imaging the differences between YFP intensity in Ala10 and Ala16 were too high for analysis with a single threshold. Therefore, two thresholds were applied, a low threshold that matched the Ala10 puncta and a high threshold that matched the Ala16 puncta but did not detect puncta in Ala10 (Fig. 2A). Importantly, the selected threshold was consistent across all images. The number of puncta per nucleus was on average 6-times higher in Ala16 than in Ala10 at the low threshold (Fig. 2B). The puncta intensity showed a similar distribution in the range 1.3-1.85 (log2) intensity for Ala10 and Ala16, and this range was considered for comparative analysis (Fig. 2C). A significant linear correlation was found between intensity and area for both Ala10 and Ala16 puncta (Fig. 2D). This positive correlation was unchanged when using the Ala16 high threshold (Fig. 2D). In contrast, the correlation between area and circularity was found to be significant only for Ala16, in both low and high thresholds (Fig. 2E). In Ala16, larger puncta lost their circularity, whereas in Ala10 was circularity remained unchanged. These differences suggest differences in aggregation between Ala10 and Ala16.

Next, we assessed aggregation in living cells using spinning disk imaging. To explore differences in puncta dynamics between Ala10 and Ala16, we imaged the muscle cells 16 hours after Dox treatment (Movie S1). For comparative kinetic analysis, single nuclei with similar YFP intensity levels were selected and three sequential timeframes (2 seconds apart) were false colored in red, green or blue, an overlay was made to assess dynamics (Fig. 2F). The puncta visible in white indicate an overlap between the three timeframes indicating a minimal puncta movement (Fig. 2F). In contrast, the non-overlayed puncta were observed as separated colors, indicating mobile puncta (Fig. 2F). In Ala10 the proportional area of white puncta was larger than in Ala16 (Fig. 2G). This suggests that, prior to aggregation, puncta dynamics in Ala10 is lower than in Ala16.

We then assessed aggregation kinetics in differentiating cell cultures that were treated with Dox 12- and 24-hour intervals (Fig. 3A). Assessment of the YFP signal in fused cells after 72 hours of Dox treatment showed remarkably less fluorescence in Ala10 compared to Ala16 (Fig. 3B). The number of puncta per nucleus was significantly lower in Ala10 compared to Ala16 (Fig 3C). Similar to the analysis in unfused cells (Fig. 2E), a negative correlation was found between puncta area and circularity in myonuclei of Ala16 fused cells (Fig. 3D). In Ala10 fused cells a correlation between puncta area and circularity was also significant, but the distribution of puncta area in Ala16 was 8-fold larger than in Ala10 (Fig. 3D). High variation between myonuclei in the same fused cell was found, moreover in Ala16 fused cells (Fig. 3B). Therefore, to visualize puncta accumulation overtime in in fused cells, we applied one YFP contrast stretch factor in Ala16 images and a 3-folds higher contrast stretch factor in Ala10 images, the same contrast stretch factor was applied to all Ala16 or Ala10 images. In the Ala10 cell cultures the mono nucleated cells were much brighter than the multinucleated cells(Fig. 3E and Fig. S4). In the Ala16 cell cultures, the difference in YFP fluorescence intensity between mono and multinucleated cells was moderate compared with the Ala10 cell culture (Fig. 3E). In the Ala10 fused cells YFP intensity remained similar over time, but in the Ala16 fused cells it increased (Fig. 3E). This visual assessment suggests that aggregation differs between fused and unfused cells and that in fused cells aggregation of Ala10 is minor compared with Ala16.

**Figure 3.**
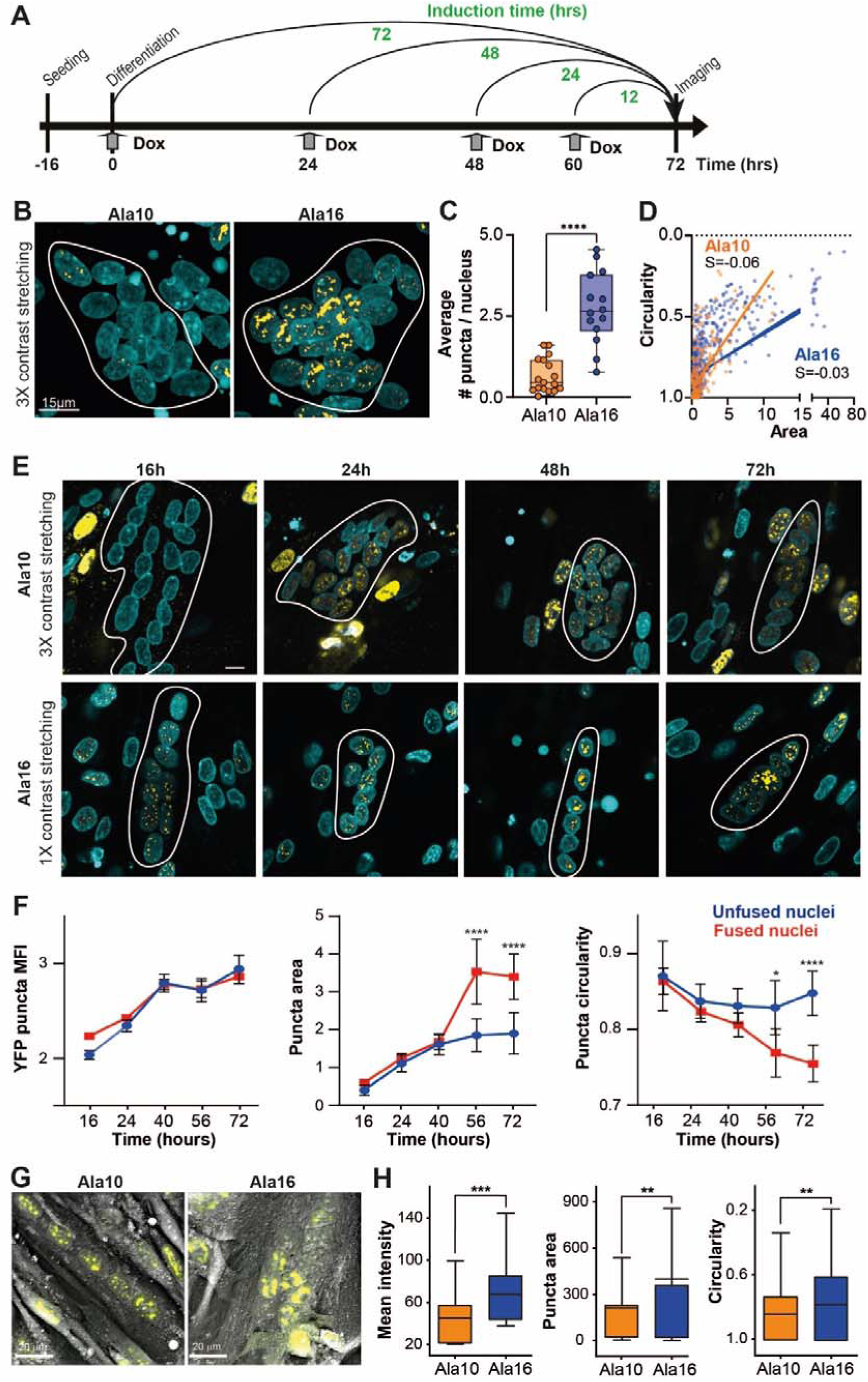
Kinetics and structure of Ala10 and Ala16 puncta in fused cells. **A** A graphical summary of the experiment to study PABPN1-puncta accumulation if fused cells. **B.** A representative image of YFP flourescence in Ala10 and Ala16 fused myotube, 72 hours after Dox induction. Equal stretching was applied for YFP signal in Ala10 and ala16. Myonuclei in fused cells are encircled. **C.** A boxplot of the average puncta number per nucleus and a violin plot of YFP MFI and in Ala10 (orange) or Ala16 (blue) fused cells. mean is from N=226 Ala10 and N=195 Ala16 nuclei for the box plot and N=147 Ala10 and N=511 Ala16 YFP puncta for the violin plot. Statistical significance was determined with one-way ANOVA for the boxplot and a Brown-Forsythe and Welch ANOVA test for the violin plot. **D.** Scatter plots of puncta area vs. Circularity. The linear regression line is depicted per genotype, and the significant slope (S) is denoted. **E.** Representative images of puncta acummulation in Ala10 and Ala16 fused cells from 12 hours till 72 hours after Dox treatment. Per genotype, the contrast stretch factor was determined on the 72 hr image and remained in all time points. The contrast strech factor in Ala1’0 was 3folds higher than in Ala16. Myonuclei in fused cells are outlined. **F.** Quantification of Ala16 YFP puncta MFI, area and circularity over 72 hours in unfused (blue) and fused (red) myonuclei. Average and standard deviation are from N=41-73 unfused myonuclei and N=63-137 fused myonuclei. **G.** Representative refractive index images of Ala10-YFP and Ala16-YFP differentiated cells, overlayed with fluorescence signal. Scale bar is 20 μm. **H.** A boxplots of YFP mean intensity (left), puncta area (middle) and circularity (right). Statistical significance was assessed with one-way ANOVA with Tukey. P<0.05; <0.01; <0.005; or <0.0001 is denoted with *; **; ***; ****, respectively.

To investigate the differences in aggregation between fused and unfused cells over time we focused on differentiating Ala16 cell cultures. We separated myonuclei from fused and unfused cells in the same plate and applied the same puncta mask on all nuclei (Fig. S5). Puncta intensity increased with time but did not differ between fused and unfused nuclei (Fig. 3F). In contrast, from 40 to 72 hours in differentiation conditions the area and circularity differed between myonuclei from unfused and fused cells (Fig. 3F). An increase in puncta area and a decrease in puncta circularity (Fig. 3F) suggests greater aggregation in fused cells compared with unfused cells.

Since the quantification of puncta could be influenced by imaging and image processing, we verified the differences in puncta structure features in fused cells using refractive index (RI) imaging combined with a fluorescence imaging platform, as implemented in NanoLive holo-tomographic microscopy. This enabled us to simultaneously acquire morphological and molecular density information, which allows for a more detailed analysis of how puncta formation can affect the structural features of subcellular compartments in fused cells. Live imaging of Ala10 and Ala16 differentiated cells revealed that in the Ala16 cells puncta intensity was higher, larger, and less circular as compared with Ala10 cells (Fig. 3G and Fig. 3H). Overall, the results from the combined RI/Fluorescence imaging corresponds with the results obtained from confocal imaging. Together, in fused Ala16 cells nuclei are more densely packed with protein aggregates than in Ala10 cells. This consistency across different imaging modalities and analytical techniques enhances the robustness of our observations.

Next, we employed transmission electron microscopy (TEM) to characterize PABPN1 aggregates in fused cells at nanometer-scale resolution. A stitched image of a fused cell created with an in-house software (Faas, Avramut et al. 2012), allowed for detailed analysis of the entire fused cell (Fig 4A). In the Dox-treated cell cultures we noticed electron-dense structures varied in morphology. These structures, which were absent in the vehicle control myonuclei, showed different electron-density from nucleoli (Fig. 4B). We classified the myonuclei based on the morphology of these structures; the smaller structures we referred to as ‘punctate’ and the larger structures as ‘bouquet’ (Fig. 4B). The bouquet appeared as encompassing the entire nucleus area or only part (Fig. 4B). To confirm that these novel structures corresponded to PABPN1-YFP aggregates, we used correlative light and electron microscopy (CLEM). By using the YFP label, we were able to correlate and overlay images of the same area acquired using fluorescence microscopy and electron microscopy. This CLEM data showed that each electron-dense aggregate corresponded to a YFP signal (Fig. 4C, encircled in pink). Some YFP signal did not correspond to an electron dense structure, due to the aggregate being above or below the 90 nm-thick section imaged in the electron microscope (Fig. 4C, encircled in orange).

**Figure 4:**
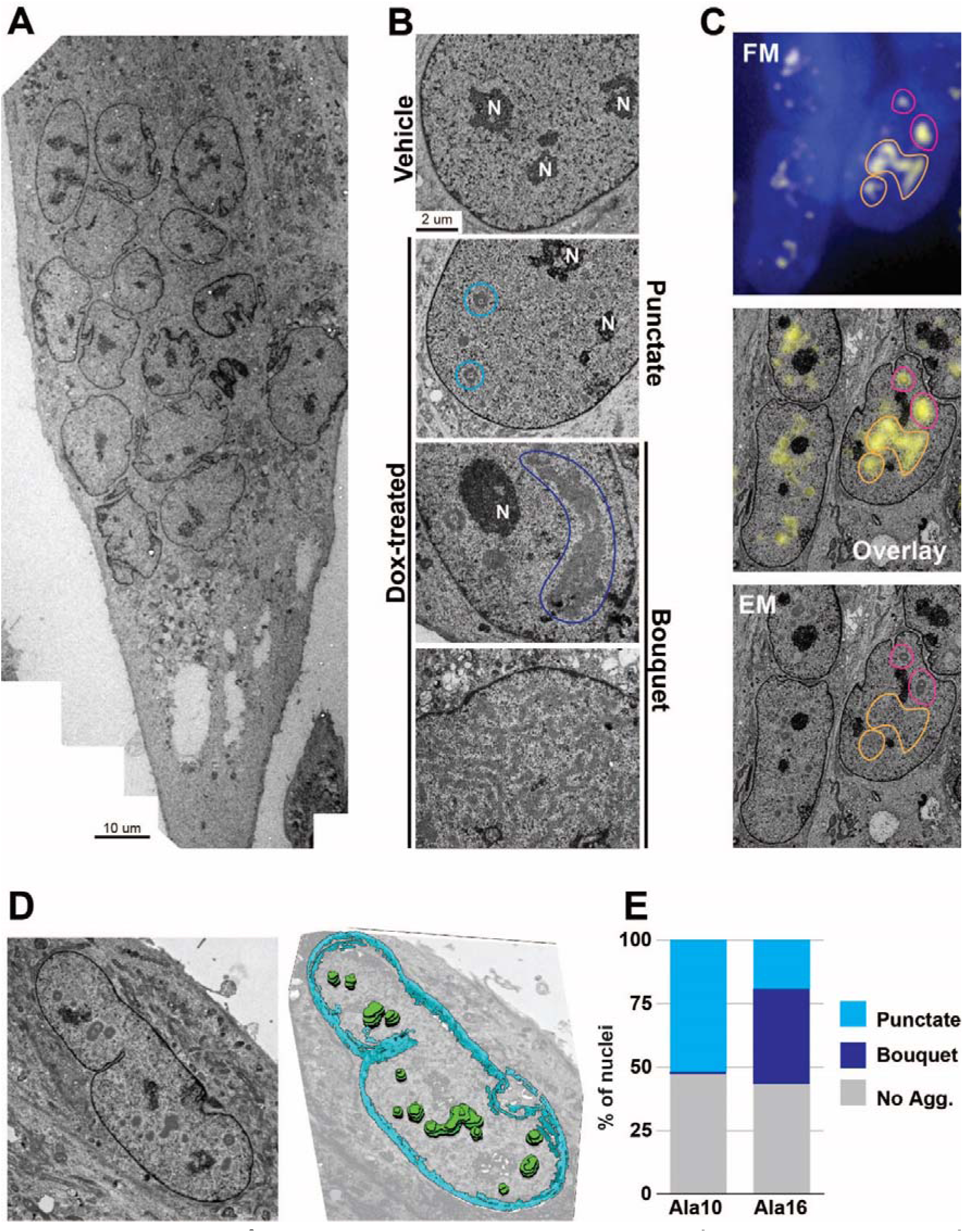
Detection of PABPN1 aggregates structures using Electron microscopy and correlative light and electron microscopy (CLEM). Images are from fused cells. **A.** EM image of a myotube in Ala16-YFP cell culture,. **B.** From top to bottom: an EM image of a nucleus in vehicle cell culture followed by aggregated structures: punctate (encircled in cyan) and two bouquet images (encircled in blue) that differ in size and shape. Nucleoli (N) are annotated. **C.** Light microscopical (FM) and electron microscopical (EM) images and overlay of the same cell area YFP signal correlating with protein aggregates structures are encircled in pink, and YFP signal without a correlation is encircled in orange. **D.** 3D reconstruction of PABPN1 aggregates in a Ala16 cell. Left, a middle section; Right, a 3D reconstruction based on 8 serial sections. Aggregates are green; the nuclear envelope is in cyan. **E.** Bar chart shows the percentage of nuclear structure (punctate or bouquet) or nuclei without aggregates in Ala10 and Ala16 myonuclei. Per genotype, 100 nuclei were sampled.

Some punctate aggregates displayed a toroidal architecture (Fig. 4B, cyan circles). Sectioning through 3D spherical, cylindrical, or toroidal structures can yield a torus in the resulting 2D slice. To determine the morphology of these aggregates, we imaged 8 sequential 200 nm-thick sections using TEM for 3D reconstruction (Figs. 4D and S6). This revealed that the toroidal structures represent hollow spheres of aggregated material, and these spheres can further aggregate into larger structures.

To assess structural differences between Ala10 and Ala16, the aggregate structures were manually assessed in 90 nm-thick sections from 100 nuclei (Fig.4E). This assessment showed that a ‘punctate’ morphology is present in 50% of the nuclei in both Ala10 and Ala16 myonuclei, whereas a ‘bouquet’ morphology was found in only one Ala16 myonucleus (Fig. 4E). From the myonuclei with electron-dense structures in Ala16, two-thirds showed a ‘bouquet’ morphology (Fig. 4E). Altogether, the EM imaging confirms the structural differences, in both area and circularity, between Ala10 and Ala16 aggregates in fused cells that was indicated from the fluorescence imaging.

### Difference in morphology and size of the aggregates affects the cell nucleus as revealed by RI imaging

Our fluorescence and electron microscopy analysis highlighted significant structural differences between Ala10 and Ala16 nuclear aggregates. To determine if the accumulation of nuclear aggregates affects nuclear architecture, we used label-free RI imaging, which allows textural features referring to myonuclei structure to be extracted. We imaged myonuclei in fused cells (Fig. 5A), and calculated entropy and Inverse Difference Moment (IDM). Entropy measures texture complexity, with higher values indicating more detailed information, while IDM evaluates image homogeneity, where higher values indicate a more uniform texture (Pantic, Dimitrijevic et al. 2016). Ala10 myonuclei exhibited higher entropy and lower IDM values compared to Ala16 (Fig. 5B). The Ala10 values were more similar to the vehicle myonuclei (Fig. 5B). While our approach does not uncover the chemical nature or the mechanism behind the structural changes, the significant difference in nuclear texture measures between Ala10 and Al16 suggests that the pathogenic aggregates impact nuclear organization.

**Figure 5:**
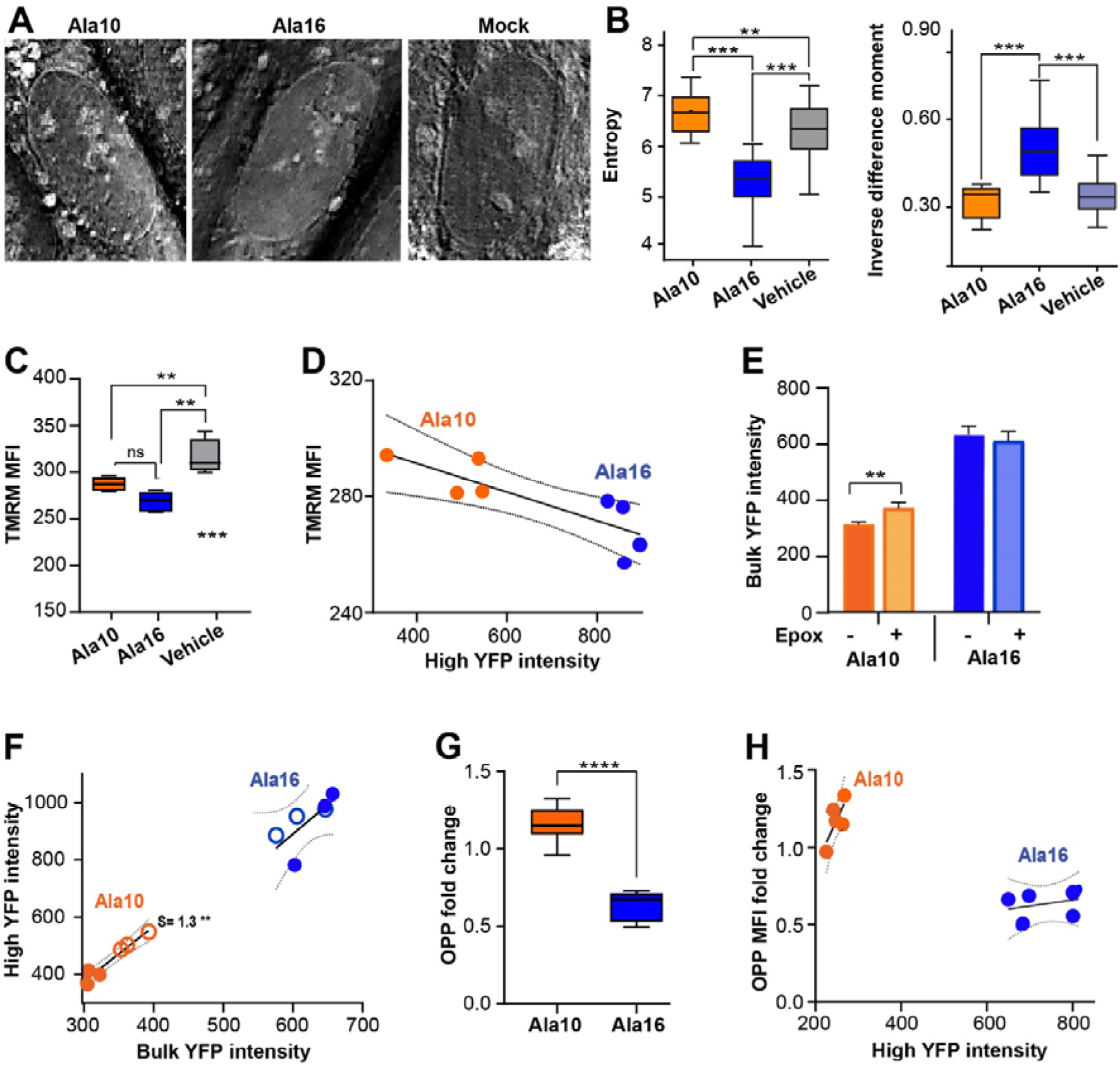
Assessment of textural and cellular differences between Ala10 and Ala16 differentiating cell cultures. **A-B** RI imaging, **A.** Representative nuclei. **B.** Boxplots of entropy or inverse difference moment in Ala10, Ala16 or vehicle expressing cells. **C-H.** HTC imaging. **C-D.** TMRM staining of mitochondrial activity. **C.** Boxplot shows TMRM mean fluorescence intensity (MFI) in Ala10, Ala16 or vehicle expressing cell cultures. **D.** A correlation plot between high YFP intensity and TMRM MFI. The regression line and 95% confidence are depicted. **E-F.** Proteasome inhibition assay in differentiation conditions followed by 6 hours epoxomicin or mock treatment. **E.** bulk YFP intensity in mock- or epoxomicin- (Epox) treated cell cultures. **F.** A correlation plot between high YFP intensity. A linear correlation and 95% confidence are depicted, a significant slope (S) is depicted. G-H. Translation efficiency in fused cell cultures assessed by OPP-Cy5 incorporation. **G.** Boxplot of OPP MFI cell cultures. MFI values show the fraction from vehicle cell cultures. **H.** A correlation plot between high YFP intensity and OPP MFI. The regression line and 95% confidence are depicted Means and standard deviations are from N=4 for C-H. Statistical significance was assessed with an ANOVA test. P<0.01; <0.005 or <0.0001 is denoted with **, *** and ****, respectively.

*Molecular and cellular mechanisms are differentially correlated with Ala10 and Ala16 PABPN1 aggregates*.

The multimodal imaging techniques employed above revealed that Ala16 aggregates are significantly larger than Ala10 aggregates. Since puncta intensity positively correlated with area in both cultures, we investigated whether puncta intensity correlates with cellular mechanisms that are regulated by PABPN1. Mitochondrial activity is reduced in OPMD (Chartier, Klein et al. 2015, Doki, Yamashita et al. 2019) and is negatively affected by reduced PABPN1 levels (Anvar, Raz et al. 2013). We measured mitochondrial activity in cells incubated with TMRM, which is sequestered in active mitochondria. Compared with vehicle cell cultures, in both Ala10 and Ala16 cell cultures TMRM intensity was significantly reduced, but TMRM intensity did not differ between wild-type and mutant PABPN1 (Fig. 5C). TMRM intensity was higher in differentiating cell cultures compared with proliferating cell cultures (Fig. S7), but cell culture conditions did not lead to a difference between Ala10 and Ala16 cell cultures (Fig. S7). A negative correlation was found between high YFP intensity and TMRM intensity but did not differ between Ala10 and Ala16 (Fig. 5D). This suggests that PABPN1 aggregation negatively impacts mitochondrial activity, regardless of aggregate size or differentiation condition.

PABPN1 protein accumulation is regulated by the ubiquitin proteosome system (Raz, Buijze et al. 2014), therefore we assessed the effect of proteasome inhibition on aggregate size. Proteasome inhibition by epoxomicin treatment in differentiated cells resulted in a higher Ala10-YFP fluorescence but Ala16-YFP intensity was unchanged (Fig. 5E). This agrees with previous observations in a mouse cell model for OPMD (Raz, Routledge et al. 2011, Raz, Buijze et al. 2014). The high YFP intensity was elevated after epoxomicin treatment in both Ala10 and Ala16, but a correlation between YFP intensity and high YFP intensity was significant only in Ala10 cells (Fig. 5F), which indicates that Ala10 aggregates are more impacted by proteasome activity compared with Ala16.

Translation efficiency is also impacted by PABPN1 levels (Mei, Boom et al. 2022). Measured by OPP-cy5 that marks active translation, translation efficiency in Ala16 differentiating cell cultures was significantly lower compared with Ala10 or vehicle cell cultures (Fig. 5G). Translation efficiency in Ala10 cell cultures was indifferent from vehicle cell culture (Fig. 5G). Moreover, in differentiating cell cultures, translation efficiency was lower in fused cells compared with unfused cells (Fig S7), further indicating the relevance of cell differentiation condition on aggregation. The correlation between high YFP intensity and translation efficiency (OPP intensity) differed between Ala16 and Ala10 (Fig. 5H), indicating that Ala16 aggregates negatively impacts translation efficiency.

In OPMD, mRNA is sequestered in nuclear aggregates (Calado, Tomé et al. 2000). In our cell model, oligo-dT-cy5 colocalized with YFP signal (Fig. 6A), and showed a nuclear sequestering as compared to vehicle cell cultures (Fig. S7E). The intensity of nuclear oligo-dT was 3-fold higher in Ala16 compared to Ala10 in differentiating cell cultures (Fig. 6B, left), whereas in proliferating cell cultures the difference was on average only 1.7-fold (Fig. 7SF). A correlation between oligo-dT and YFP signal in nuclear area was on average 6-fold higher in differentiating Ala16 compared to Ala10 (Fig. 6B, middle), but in proliferating cell cultures the difference was on average 2.8-fold higher (Fig. 7SF). This demonstrated mRNA entrapment in Ala16 aggregates is higher than in Ala10. Importantly, mRNA entrapment is affected by cell differentiation. We then assessed mRNA nuclear export by the nuclear to perinuclear oligo-dT intensity ratio. The ratio was also higher in Ala16 (Fig. 7B, right) supporting the notion that nuclear entrapment of mRNA impacts mRNA nuclear export. We also determined whether nuclear export is affected in Ala10 and Ala16 cells using leptomycin B (LMB), a nuclear export inhibitor. LMB treatment impacted the nuclear to cytoplasmic oligo-dT ratio in Ala10 proliferating cell cultures but not in Ala16 cell cultures (Fig. S7H). This implies that nuclear export is malfunctioning in Ala16 cells. LMB treatment in differentiating cell cultures did not impact oligo-dT subcellular accumulation (Fig. S7H), signifying that the regulation of protein nuclear export differs between cell culture conditions.

**Fig. 6:**
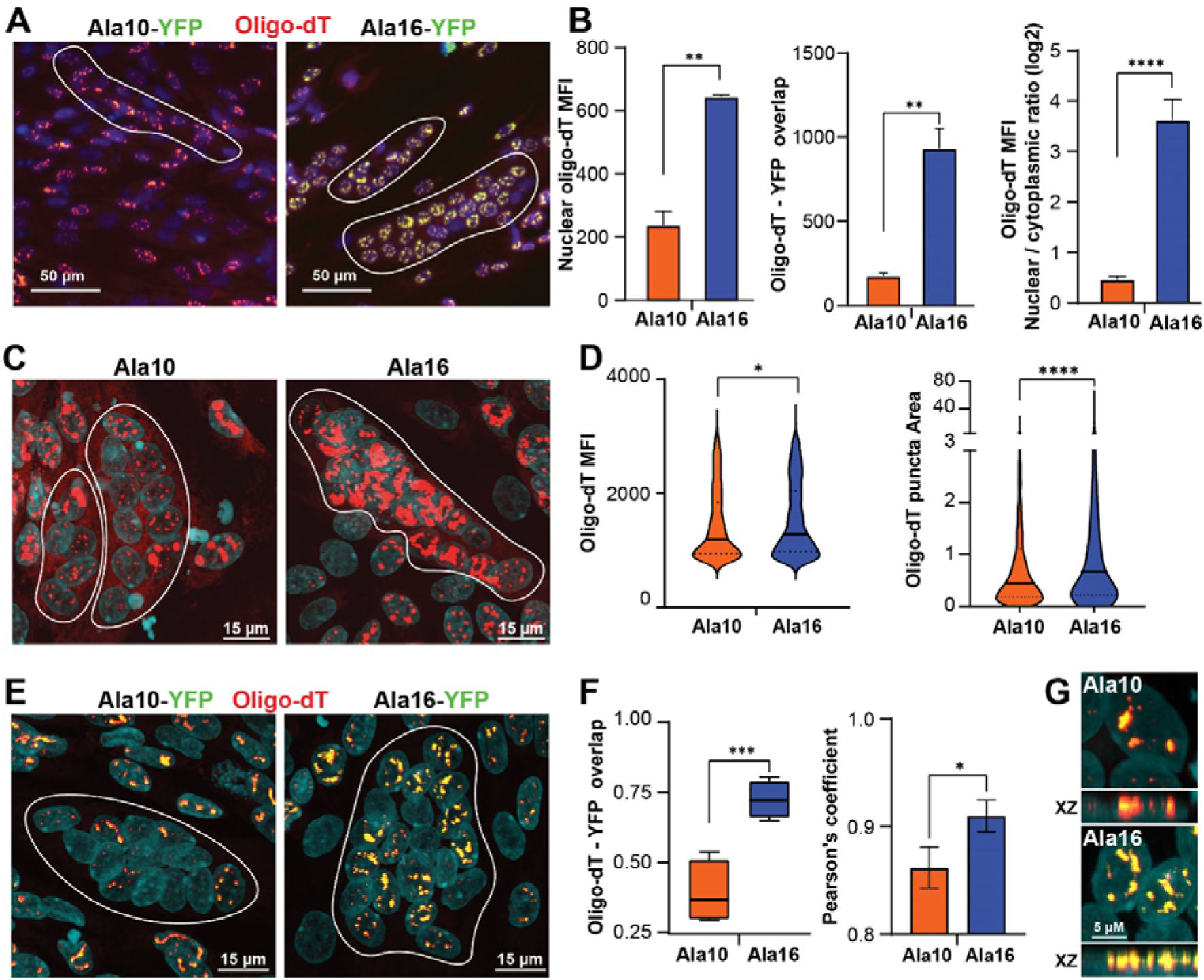
mRNA binding properties of Ala10 and Ala16 aggregates. **A.** Representative images of Ala10 and Ala16 fused cells labeled with Oligo-dt-Cy5. **B.** Bar graphs show differences in Oligo-dT MFI, nuclear to cytoplasmic ratio, and Oligo-dT overlapping YFP signal between Ala10 and Ala16 myonuclei. Statistical significance was assessed by ANOVA test. **C.** Confocal images of fused cells labeled with oligo-dT-Cy5. Fused cells are encircled with a white line. Nuclei are counterstained with DAPI, the scale bar is 15 μm. **D.** Violin plots show oligo-dT puncta MFI and area. Statistical significance was assessed with an unpaired t-test. **E.** Representative confocal images of YFP (green) and oligo-dT (red) in Ala10 and Ala16 cells. Yellow denotes an overlap between red and green. Fused cells are encircled with a white line. Scale bar is 15μm. **F.** Boxplot of overlap values YFP with oligo-dT and oligo-dT with YFP. Statistical significance was assessed with an ANOVA test. Bar charts show the Pearson correlation between YFP and oligo-dT. P<0.05; <0.01; <0.005 or <0.0001 is denoted with *, **, ***, ****, respectively. **G.** An XY and XZ plane of a representative 3D image in Ala10 and Ala16 myonuclei. YFP is in green, oligo-dT in red and yellow denotes an overlap. Scale bar is 5 µM.

Finally, we verified the results obtained by the HCS imaging using confocal imaging and analysis in fused cells (Fig. 7C). Similar to YFP intensity, in Ala10 cell culture, the oligo-dT signal in myonuclei in fused cells was lower than in unfused cells (Fig. 7C). However, in Ala16 the oligo-dT signal in myonuclei in fused cells was higher than in unfused cells (Fig. 7C). Oligo-dT puncta intensity and area were higher in Ala16 compared to Ala10 (Fig 7D). A colocalization between oligo-dT and YFP was calculated from images with both fluorophores (Fig. 7E). Per pixel, the oligo-dT overlap with YFP puncta was significantly higher in Ala16 compared with Ala10 (Fig. 7F, left), and the Pearson correlation was significantly higher in Ala16 (Fig. 7F, right). We then assessed the colocalization between YFP and oligo-dT in 3D confocal images of myonuclei. In Ala16 myonuclei the overlap between YFP and oligo-dT was nearly complete, but in Ala10 part of the oligo-dT signal did not overlap with YFP fluorescence (Fig. 7G). Thus, a 3D visual assessment matched with the overlap statistical analysis. Moreover, the overlap and correlation scores calculated from confocal images are consistent with HCS imaging demonstrating that YFP aggregates co-localization with oligo-dT signal is significantly higher in Ala16.

## Discussion

Our analysis, supported by advanced microscopy across five distinct imaging modalities, uncovered aggregation at the micro- to nano-scale range, and revealed differences between pathogenic and non-pathogenic nuclear aggregates. At the nanoscale, the pathogenic aggregates are characterized by their ‘bouquet’ formation, composed of connected punctate units, whereas the non-pathogenic form are non-connected punctae (Fig. 4). At the microscale, the pathogenic aggregates exhibit high fluorescence intensity, larger size, and reduced circularity as compared with the aggregates of the non-pathogenic protein (Fig. 3). The distinction between non-pathogenic and pathogenic PABPN1 aggregates was more pronounced in differentiated multinucleated cells compared with non-differentiated cells (Fig. 3). Previous studies in non-muscle proliferating cell cultures reported only limited differences between aggregates of the wild type and the expanded PABPN1 aggregates (Raz, Abraham et al. 2011, Guan, Jiang et al. 2023). In non-muscle proliferating cells, slow protein dynamics of the expanded PABPN1 have been suggested to correlate with higher aggregation (Raz, Abraham et al. 2011, Guan, Jiang et al. 2023). Our time-lapse imaging in differentiating cells suggests that the single punctae of the wild-type PABPN1 mobility is higher than the single punctae of the expanded PABPN1, but when puncta accumulate, their mobility within the wild-type PABPN1 is reduced compared with the expanded PABPN1. Together, this suggests that the self-assembly of wild-type and pathogenic PABPN1 differ, potentially due to interactions with other nuclear proteins or molecules. Protein mobility is affected by multiple factors, but in general, the larger the molecule or complex is, the slower the mobility will be (Ma, Wan et al. 2020). Aggregates of wild type and pathogenic PABPN1 are of nearly similar size, but their differences in aggregation suggest differences in nuclear protein interactors, such as other RNA-binding proteins (Gavish-Izakson, Velpula et al. 2018) or nuclear chaperones’ regulators of protein folding (Vonk, Rainbolt et al. 2020). In addition, irreversible mRNA binding to the RNA binding domain, as we show here for the aggregated PABPN1, could also impact protein mobility (Jankowsky and Harris 2015).

Most disease-associated protein aggregates are presented in post-mitotic cells, such as neuronal cells or fused muscle cells (Shastry 2003). RNA-protein interaction patterns are impacted during cell differentiation (Trendel, Schwarzl et al. 2019). Protein-protein interactions are also changed during cell differentiation. Specifically, the chaperone network system is changed in differentiated neuronal cells (Vonk, Rainbolt et al. 2020), underlying differences in the maintenance of proteostasis between proliferating and differentiated cells. Our experiments in differentiated muscle cells highlight the importance of studying nuclear aggregates in cell models that mimic disease conditions. and on disease mechanisms In our cell-based analysis, we explored a correlation between cellular function and PABPN1 aggregates. Decreased mitochondrial activity is a key factor in NMDs (Shastry 2003, Connolly, Theurey et al. 2018), including in aging muscles and in OPMD (Anvar, Raz et al. 2013, Chartier, Klein et al. 2015). Mitochondrial activity did not discriminate between wild type and pathogenic PABPN1 aggregates. Inhibition of the proteasome discriminated between wild-type and the pathogenic PABPN1 aggregates. Using epoxomicin to inhibit the proteasome, we saw an increase in Ala10-YFP aggregate accumulation, indicating their usual degradation by the proteasome. However, the same treatment did not change Ala16-YFP, suggesting that either the Ala16 protein is blind to the proteasome, or the proteasome is malfunctioning in Ala16 cells. The impairment of the proteasome in Ala16 cell culture points to a functional significance in OPMD, agreeing with previous studies (Anvar, t Hoen et al. 2011, Ribot, Soler et al. 2022). Similarly, we show here that PABPN1 aggregates negatively affect translational efficiency, which was complemented by mRNA sequestering in PABPN1 nuclear aggregates (Fig. 5 and Fig. 6). Also, in OPMD, poly(A) RNA is sequestered in myonuclei (Calado, Tomé et al. 2000), further strengthening our cell model as a relevant disease model. Notably, oligo-dT staining showed a substantially higher nuclear to cytoplasmic poly(A) ratio in Ala16 cells and diminished mRNA nuclear export (Fig. 6) An effect on mRNA nuclear export on translation efficiency was previously reported (Katahira 2015). Our results suggest that nuclear aggregates entrap mRNA, leading to reduced mRNA nuclear export and subsequently reduced translation efficiency and impaired cellular function.

In summary, in this study we combined five distinct imaging modalities, which allowed us to explore the structural characteristics of PABPN1 aggregates in a novel cell model for OPMD and to correlate between aggregates to cellular function. Combining imaging approaches, ranging from the micro to nano scale, enabled an in-depth investigation of nuclear aggregates structure and kinetics in differentiated cells. We demonstrated structural differences between pathogenic and non-pathogenic aggregates that account for cellular changes that affect muscle function. The pathogenic aggregates are larger less circular, and more dynamic, compared with the aggregates of the non-pathogenic protein. We show a correlation between aggregate size and mRNA nuclear entrapment and translation efficiency in differentiated muscles. In addition, we showed differences in aggregates between proliferating to differentiating cells, which strengthen the need to study protein aggregation in a disease-relevant cell condition. The cell model we show here, together with the range of imaging techniques we applied, opens an opportunity to unravel the mechanisms wiring protein dynamics in nuclear aggregates in future studies.

## Acknowledgements

This study is financed by PPP Health∼Holland and argenx. We thank Danish Khan for assisting in the generation of the muscle cell model.

