## Supplemental file for "Correlating Protein Aggregate Structure with Cellular Function in Differentiated Muscle Cells: Discriminating Pathogenic from Non-Pathogenic Forms"

### Supplementary data

#### Table S1. List of used antibodies for western blotting and immunofluorescence

**A: primary antibodies**

| Protein | Dilution | | Host | Supplier | Cat. number |
|---|---|---|---|---|---|
| | IF | WB | | | |
| GAPDH | - | 1:5000 | anti-Mouse | Invitrogen | MA5-15738 |
| PABPN1 | - | 1:4000 | anti-Rabbit | Homemade | |
| Histon2B (H2B) | - | 1:1000 | anti-Rabbit | Proteintech | 15857-1-AP |
| MyHC | 1:250 | - | anti-Mouse | DSHB | MF20 |

**B: Secondary antibodies**

| Target | Fluorophore | Dil. | Supplier |
|---|---|---|---|
| Anti-mouse | Cy5 | 1:5000 | Thermo Fisher Scientific |
| WB anti-Rb/M | 800/680 | 1:10000 | IRDye 800CW or IRDye 680RD (LI-COR) |

#### Supplementary figures

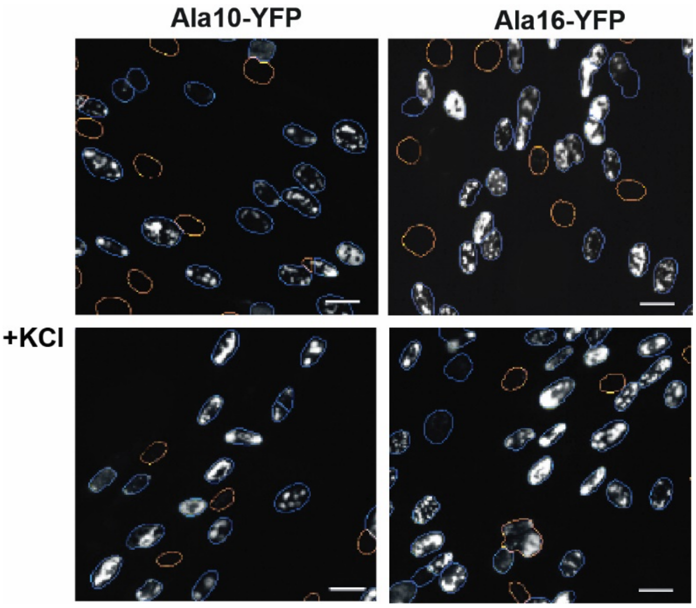


**Figure S1: Representative images of Ala10-YFP and Ala16-YFP cell culture with and without KCl treatment.** In orange are highlighted nuclei that were excluded from analysis, and in blue the included nuclei. The scale bar is 40 mm.

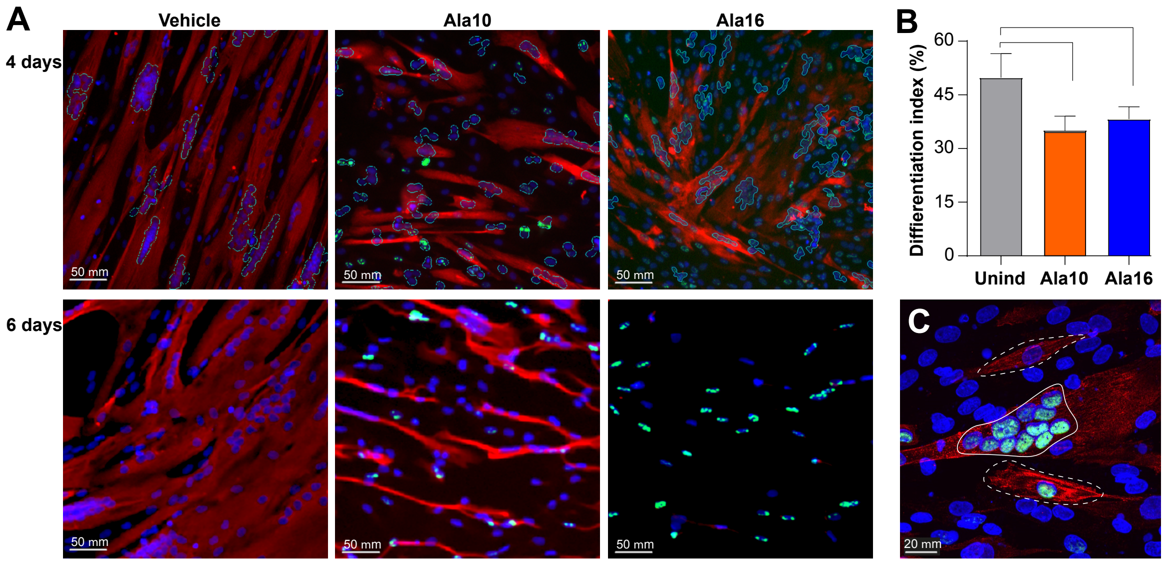


**Figure S2: Ala10 and Ala16 effect during muscle cell differentiation.**

**A.** Representative images of vehicle, Ala10-YFP and Ala165-YPF cell cultures after 4 or 6 days in differentiation condition. Dox treatment was in proliferating conditions, one day before differentiation condition. Differentiated cells are marked with MyHC (red) and multinucleated cells are highlighted by the segmentation blue line in the 4 days cultures. A longer incubation in differentiation condition (6 days) led to detachment of differentiated cells in Ala10 and Ala16 cells whereas in control cell cultures differentiated cells remained intact. In cells undergoing detachment MyHC intensity is high due to compression of the signal. In Ala16 cell detachment was more pronounced than in Ala10 indicating poor attachment after fusion. This observation agrees with previous studies showing that PABPN1 levels lead to cytoskeleton disorganization and poor cell mechanics properties in muscle cells. The scale bar is 50 μm. **B.** Differentiation index quantification in 4-days differentiating cell cultures. Average and standard deviations are from N=3. Statistical significance was assessed with a student's t-test, $p<0.05$ is denoted with *. **C.** A confocal image of Ala16 differentiated cell culture. Differentiated cells are labeled with MyHC, myonuclei in a fused cell are encircled with a continuous line. Cells expressing MyHC but unfused are encircled with a dashed line. The scale bar is 20 μm.

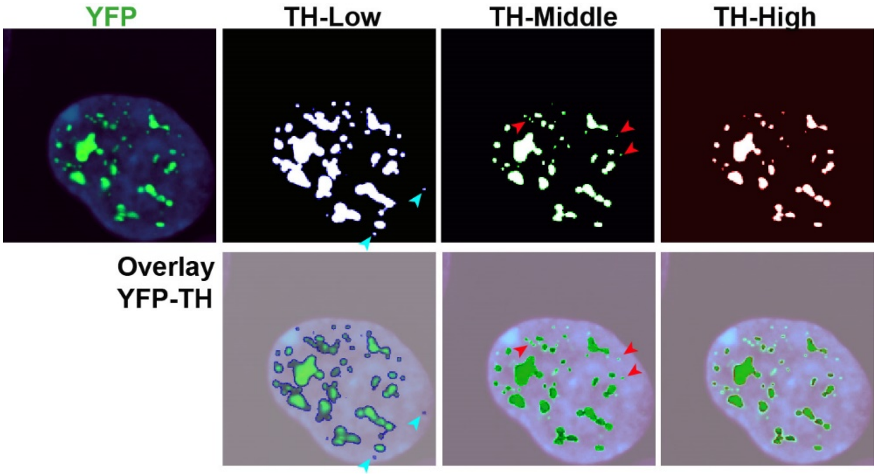

**Figure S3: Threshold criteria for puncta quantification.** Left is a fluorescence image (DAPI is in blue and YFP is green), and a mask of the YFP signal with increasing thresholds are to the right. An overlap between the YFP image and the threshold image is depicted in the lower panel. The cyan arrowheads point to puncta in TH-Low but are not detected in TH-Middle or TH-High. The red arrowheads point to puncta that are missing by TH-High. Most overlap was found between TH-Middle and TH-High.

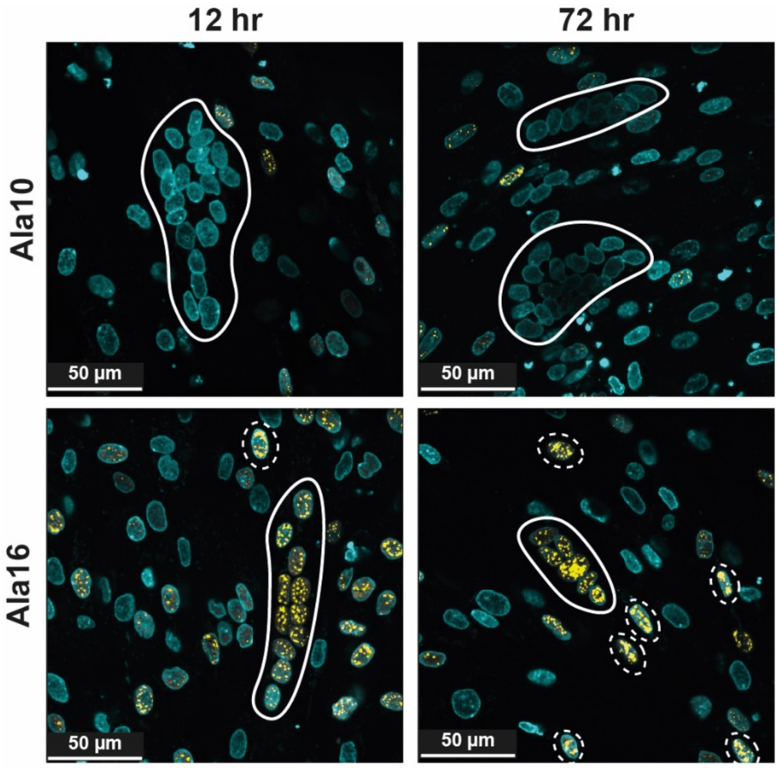


**Figure S4: PABPN1-YFP accumulation over time.** Representative images of Ala10 and Ala16 differentiated cell cultures after 12 and 72 hours of Dox treatment. Equal stretching of the YFP signal is applied to both images. Myonuclei in fused cells are encircled in continuous line. Encircled in dashed lines are nuclei in unfused cells. The scale bar is 50 µm

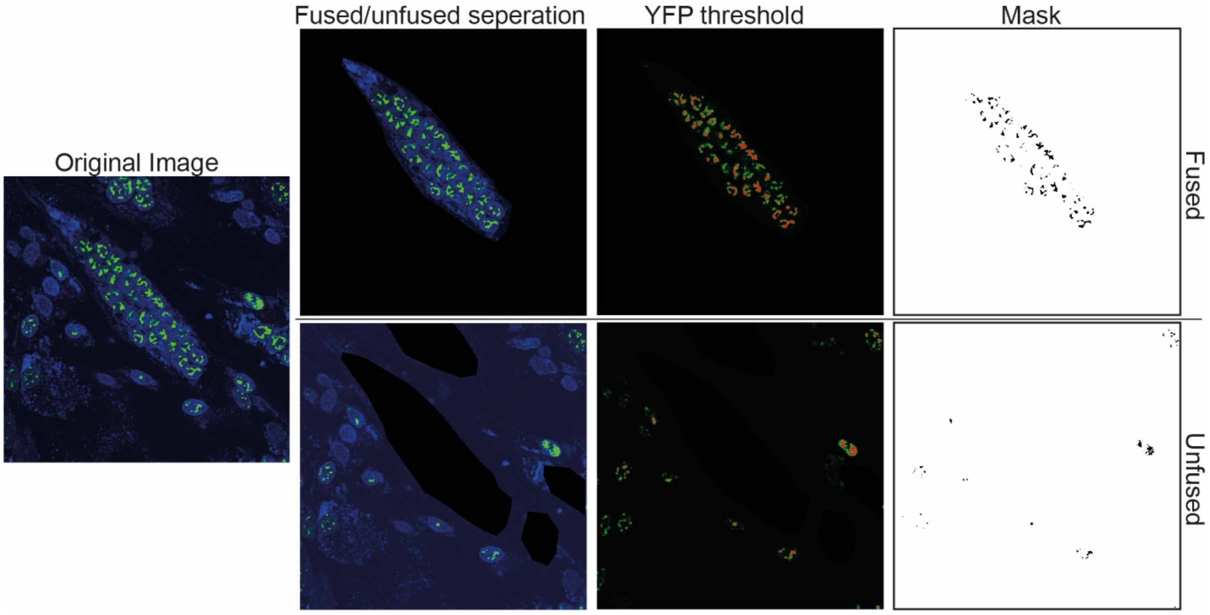


**Figure S5: Analysis of YFP puncta in unfused and fused cells from a single image.** Fused cells were cut out of the image. Subsequently, the analysis macro was run on the remaining myoblasts or on the isolate d fused cells.

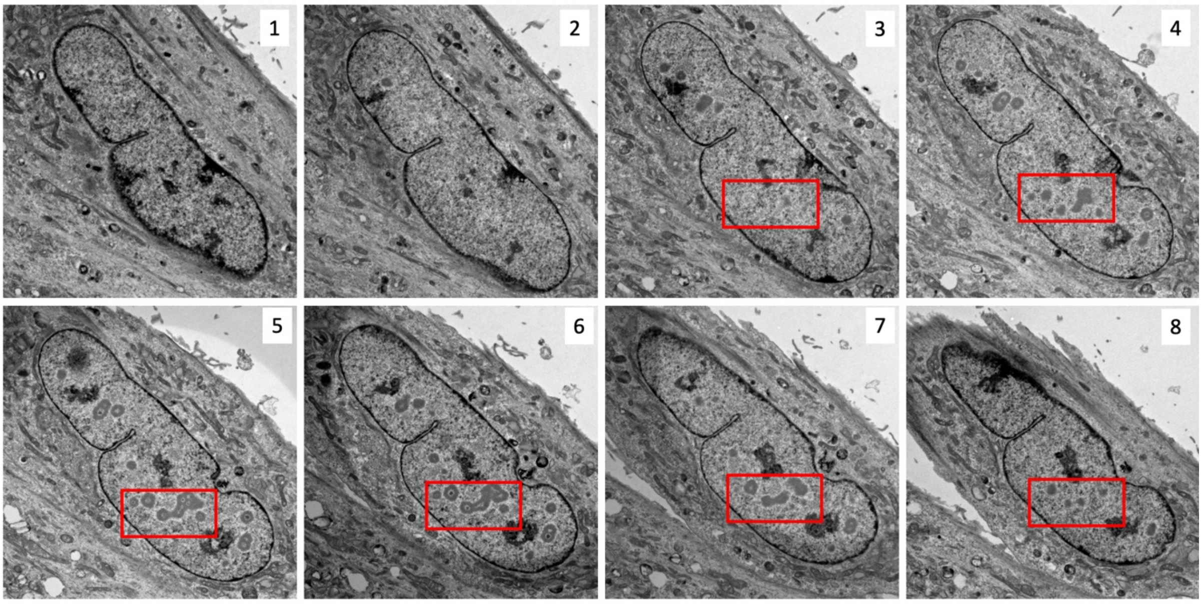


**Figure S6: Serial sections of a nucleus from an Ala16-YFP induced cell culture.** The number of each sequential serial section is depicted, section thickness is 200 nm. The same region is indicated with a red box.

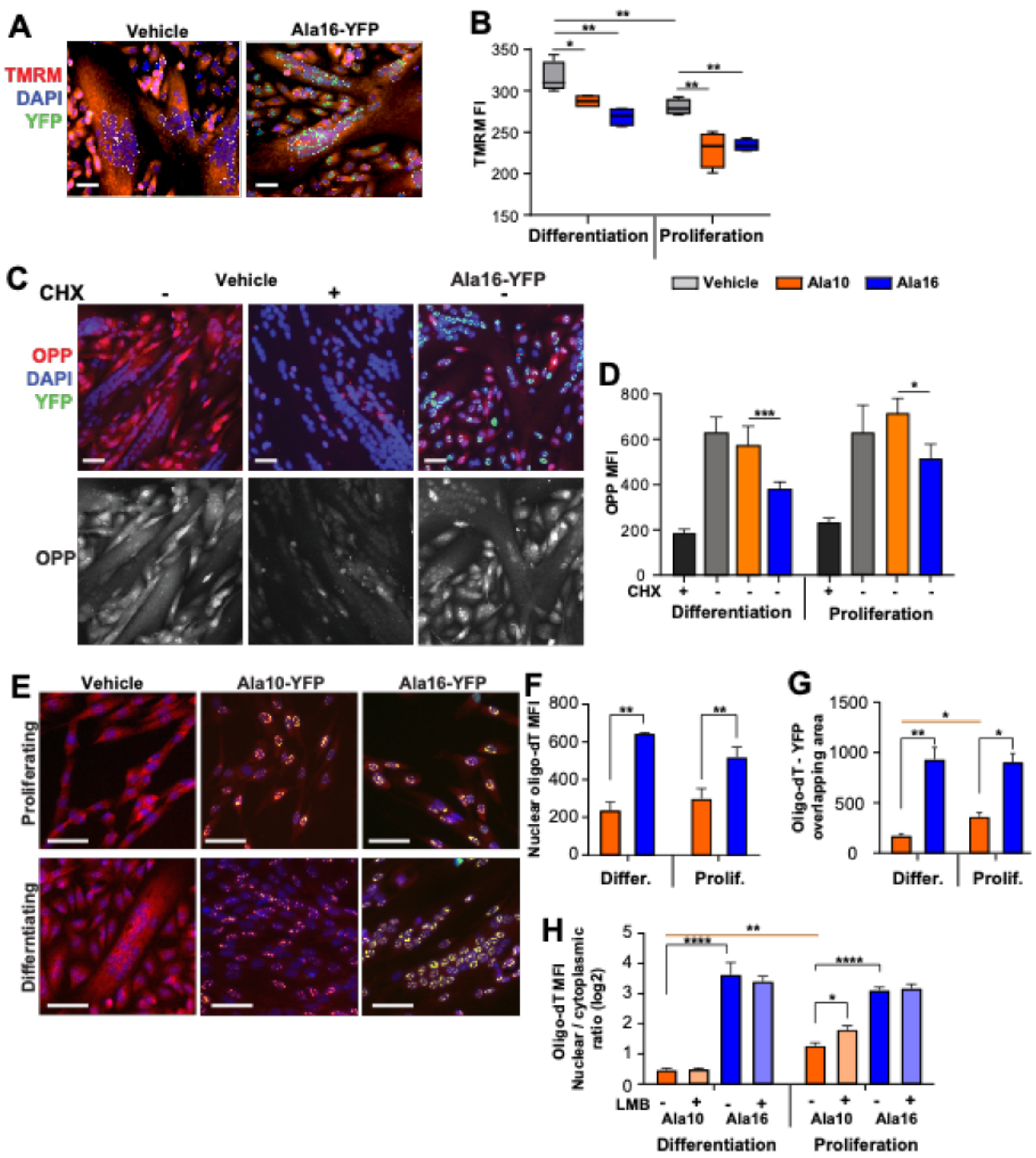


**Figure S7: Activity of cellular mechanisms in proliferating and differentiating Ala10 and Ala16 cell cultures.** Cell cultures were treated with Dox for three days. **A.** Representative images of vehicle and Ala16-YFP differentiated cell cultures incubated with TMRM (red), YFP (green). **B.** Boxplot of TMRM MFI in proliferating and differentiating cell cultures. Average and standard deviation is from N=4 biological replicates. **C.** Representative images of vehicle and Ala16-YFP differentiated cell cultures labeled OPP-555 (red). Ala16-YFP (green). Treatment with cycloheximide (CHX) is a negative control. **D.** OPP MFI in proliferating and differentiating cell cultures. Average and standard deviation is from N=4 biological replicates. **E.** Representative images of proliferating and differentiating cell culture conditions in vehicle, Ala10-YFP and Ala16-YFP cell cultures labeled with oligo-dT-Cy5 (red). YFP is in green. **F.** Nuclear oligo-dT MFI in proliferating and differentiating cell culture. **G.** YFP-Oligo-dT overlapping in proliferating and differentiating cell cultures. **H.** Nuclear to cytoplasmic Oligo-dT ratio in differentiation and proliferation cultures Mock- or LMB- treated. N=3 biological replicates. In B), D), F), G) and H) the average and standard deviations are from N=4. Statistical significance was determined with one-way Anova. A), C), and E) Scale bar is 50 μm

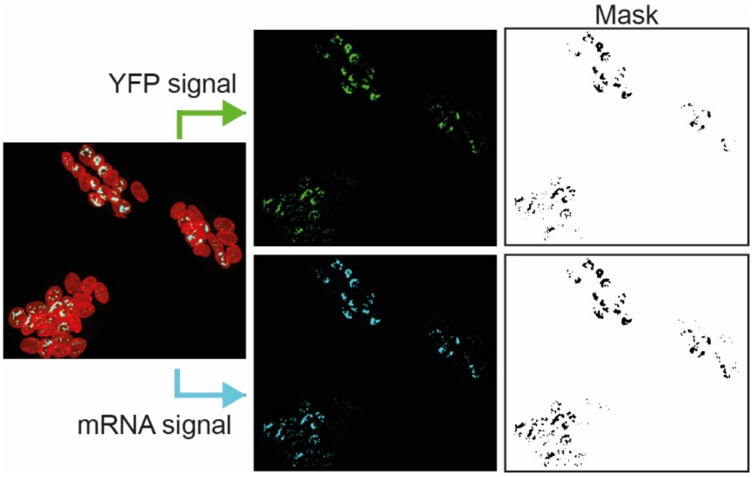


**Figure S8: Mask production of YFP and mRNA signals from images of myonuclei in fused cells for quantification.** The YFP and mRNA channels are isolated after which a mask is produced. These masks are used in the YFP-mRNA overlap quantification and YFP-puncta structure analysis.
